# Temporal dynamics of trauma memory persistence

**DOI:** 10.1101/2023.02.20.529179

**Authors:** Michael B. Bonsall, Emily A. Holmes

## Abstract

Traumatic events lead to distressing memories, but such memories are made all the worse when they intrude to mind unbidden and recurrently. Intrusive memories are a hallmark of several mental health disorders including posttraumatic stress disorder (PTSD) and can persist for years. Critically, the reduction of intrusive memories provides a treatment target. While cognitive models for psychological trauma exist, these lack formal quantitative structure and robust empirical validation. Here we develop a mechanistically-driven, quantitative framework to extend understanding of the temporal dynamic processes of trauma memory. We show how the marginal gains of interventions for intrusive memories can be enhanced as key properties of the intervention vary. Validating the framework against empirical data highlights that while emerging interventions to reduce occurrence of intrusive memories can be effective, counter-intuitively, maintaining these memories in a sufficiently reactivated state is essential for preventing their persistence.

**Author Summary:** Intrusive memories and flashbacks after trauma are prominent in several mental disorders. Quantifying these intrusions is the aim of the current study. While many conceptual models for trauma memory exist, none provide a mechanistic framework for validating experimental or clinical evidence. Our approach is to develop a probabilistic description of memory mechanisms to link to the broader goals of trauma treatment. Analysis shows how critical attributes of the framework such as intervention strength and reminder strength determine success in managing intrusive memories. Validation with empirical data shows how the framework can be parameterized and predictions evaluated against observed outcomes. In this way neural mechanisms associated with memory can be combined with broader cognitive processes.

## Introduction

Traumatic events (such as physical or sexual assaults, disasters, war experiences) are widespread [1], causing significant distress and morbidity, and a range of mental disorders. Posttraumatic stress disorder (PTSD) is characterised by ‘recurrent, involuntary and intrusive distressing memories of the traumatic event(s)’ [2]. What is special about this form of memory is that is it not only highly emotional [3], but it is thrust into mind unexpectedly against one’s will [4], and can persist for years: henceforth we referred to these as *intrusive memories*. For trauma survivors, forgetting trauma might be a long-term goal, but counterintuitively the deliberate recall of trauma memories is key in evidence-based psychological therapies [5]. One hypothesis is that under some circumstances recalling memories can temporarily return them to a malleable, labile state [6,7]. This can be achieved via a so-called ‘reminder cue’ where a simple stimulus (such as a word, a smell or a visualization) acts to reactivate memory into a labile form. Critically, during this labile period, memories may be altered/disrupted (or left uninterrupted), before reconsolidating back into long term memory [8]. The fundamental idea that consolidated memory is not permanent [9] but could again become available to alteration over a finite time window following a reminder (inferred to initiate memory reactivation) is termed ‘memory reconsolidation’ [10–12]. Memory alteration following retrieval plus various pharmacological or behavioural interventions has been achieved [13–16], though not without controversies and challenges [17]. This process suggests potential for trauma treatment innovation with procedures designed to interfere with memory reconsolidation [18,19], and critically here to make these intrusive trauma memories become non-intrusive.

While psychological models for the implications of psychological trauma are reasonably well developed [20–23], these approaches often lack quantitative predictions. Conceptual models are underpinned by the idea that a key psychopathological form of trauma recall is characterised by intrusive memories, and advances in these conceptual models have focused on developing neural bases for the combination of inflexible involuntary memories with voluntary, flexible memory [24, 25]. Relatedly, elsewhere, we have argued for a hierarchical mechacognitive framework in which neural mechanisms are embedded in cognitive processes for focal mental health symptoms [26,27].

To this end, here, together with empirical validation, we use a novel quantitative approach for investigating the temporal dynamics and persistence of intrusive memories after trauma within a memory reconsolidation framework. This framework uses probabilistic descriptors of transitions from one memory state to another. Here the processes of memory updating are described as a series of stochastic events culminating in the reconsolidation of a memory into a non-intrusive state. Our aim is to use this framework to describe how the intended reactivation of an intrusive memory (iM) via a reminder cue, followed by a behavioural task intervention can affect the probabilities of memories existing in different states. For modelling intrusive memories, our stochastic model is divided up into four distinct states (Figure 1a): (i) initial trauma; (ii) consolidated iM; (iii) reactivated iM; and (iv) nonintrusive form of memory (niM) – whereby a memory is rendered non-intrusive by the intervention.

**Figure 1.**
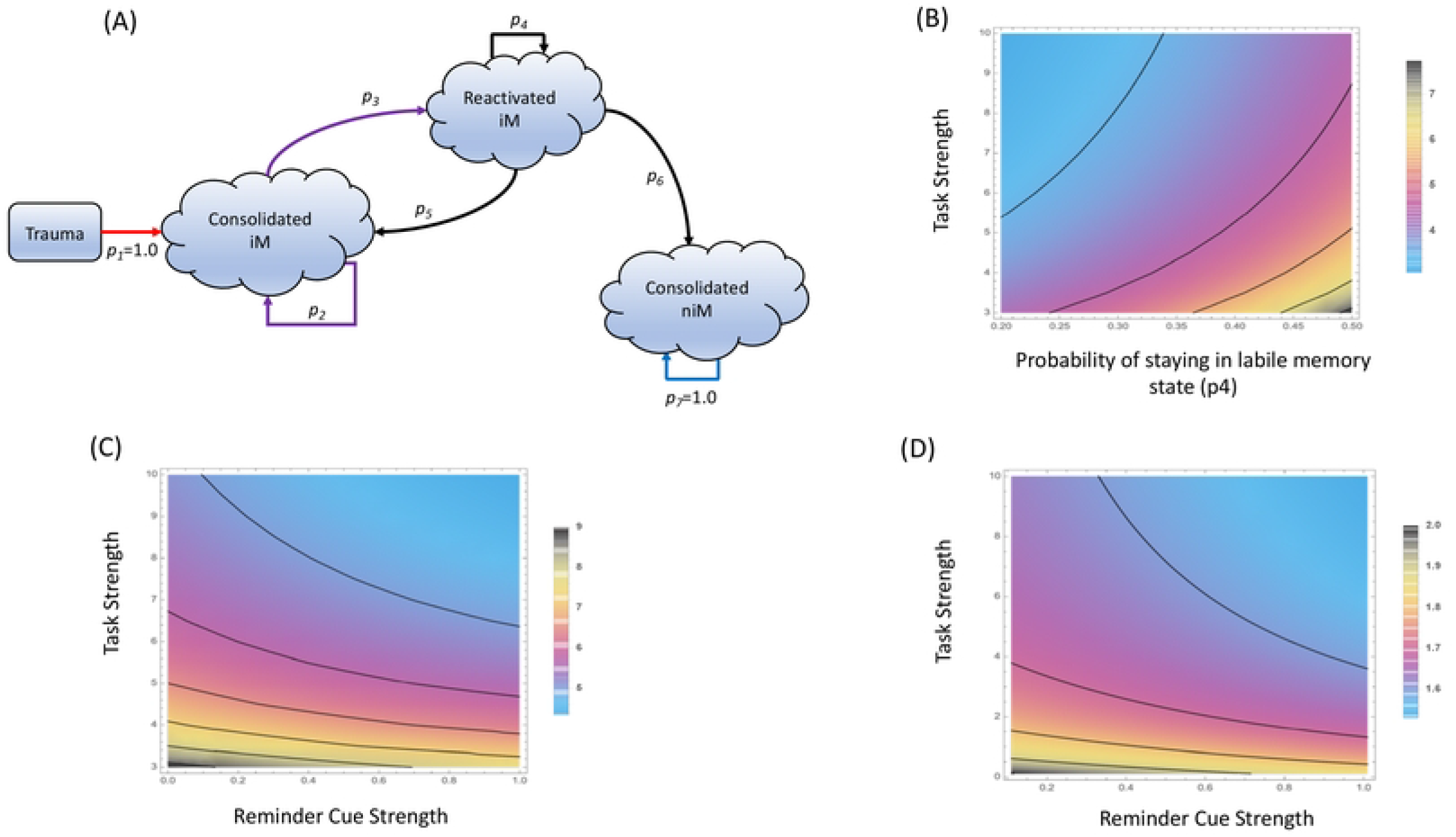
Persistence time of intrusive memories. (A) Schematic of the trauma model for different intrusive memory (iM) and non-intrusive memory (niM) states. Transitions are represented by different probabilities (p_1_ to p_7_). Coloured arrows represent different rows in the transition matrix. (B) The effects of task strength and probability of memories staying reactivated (p_4_) maintaining intrusive memories in a reactivated state on persistence of intrusive memories. Expected time in the iM state increases as task strength weakens and/or probability of staying in the reactivated state increase. Beyond certain task strength little further reduction of time in the iM state is achieved. (C) The effects of task strength and reminder cue strength on persistence of intrusive memories such that different combinations of task strength and reminder cue strengths minimize time in the iM state. (D) The effects of task strength and reminder cue strength on mixing times before memories absorb in the non-intrusive memory (niM) state such that different combinations of task strength and reminder cue strengths minimize time for memory to consolidate into the nonintrusive state. (Colours represent time in intrusive memory state)

Importantly, we define a set of probability transitions. These are the probability that after a traumatic event a given intrusive memory consolidates; here we assume that this always occurs (so p_1_=1.0) (but this need not be the case, see [26]), the probability that an intrusive memory, when spontaneously experienced, reconsolidates unaltered (p_2_), the probability that the intrusive memory is reactivated by a reminder cue (p_3_), the probability that memory stays in a reactivated state (allowing a time window for alteration) (p_4_), the probability that a reactivated memory reconsolidates as an intrusive memory and remains unaltered or is even strengthened (p_5_), the probability that the reactivated memory reconsolidates as a non-intrusive form of memory which is altered and weakened by the treatment intervention (p_6_) and the probability that the non-intrusive form of memory remains consolidated (so p_7_=1.0).

Critical to understanding how a trauma memory can be rendered non-intrusive is (i) that the intrusive memory can be reactivated with a reminder cue (p_3_) and, (ii) that a task intervention can determine whether an intrusive memory reconsolidates in an altered form or not (p_5_). With this framework, it is then feasible to determine measures such as the expected time to absorption into the non-intrusive memory (niM) state, the expected intensity and the number of visits to the reactivated iM state before absorption into the niM state - all as a function of the task intervention, and/or the reminder cue.

## Results

### Modelling Intrusive Memory Dynamics

Analysis reveals that the expected time that memories remain in an intrusive state is dependent on the probability of maintaining a memory in the reactivated state (p_4_), and parameters associated with task strength and reminder cue strength (Figures 1b-c). Expected time in the intrusive memory state increases as task strength weakens and/or the probability of memories being in a reactivated state increase. Of key importance, is that beyond a critical level of task strength, little further reduction of time in the intrusive memory state is achievable (Figure 1b). Combinations of multiple task strengths can also minimise the time memories stay in the intrusive memory state (Figure 1c).

Mixing time analysis determines how long it takes for memories to absorb into the non-intrusive memory state. As task strength and reminder cue strengths increase, mixing times are minimised before memories enter a non-intrusive form (Figure 1d). Again, beyond critical combinations of task strength and reminder cue strength little further minimization of mixing times is achievable. An upper bound on how quickly memories reach the non-intrusive state can be derived from an inequality analysis (see Methods). This shows that the upper bound is critically determined by the probability of memories being held in the reactivated state (p_4_). High probability of memories remaining in the reactivated state can lead to long times before memories reach the absorbing state (non-intrusive memory state). The shape of this relationship suggests that there are physical limits beyond which any further balancing of memories being in the reactivated state leads to no further gains in how quickly memories move into the non-intrusive memory state (niM).

### Empirical Validation

Empirically, we can use the stochastic framework to validate our memory model with experimental and/or clinical data in which intrusive memories have been manipulated [28]. To validate our model, we use a dataset from an existing memory reactivation – reconsolidation study [29]. In this study to examine memory updating mechanisms to reduce the persistence of intrusive memories, participants viewed a film with traumatic content and recorded their intrusive memories to the film for 24 hours (allowing the initial memory consolidation to occur). A day later, participants were randomised to one of four groups: no task control, memory reminder cue with task, task only, or memory reminder cue only. The memory reminder cue was briefly viewing (2 seconds) film stills associated with specific intrusive memories. The task intervention involved playing the computer game Tetris using mental rotation to optimize the placement of coloured blocks. Participants kept diaries of the number of intrusive memories over the subsequent week. From these diaries, estimates for the unknown probabilities (p_2_ to p_6_) in the transition matrix can be determined directly from empirical parameterization, assumptions about the distribution of intrusive memories and/or regression-based approaches (SI).

Using assumptions that intrusive memories follow a discrete-valued Poisson distribution (SI), the expected probabilities can be estimated using mean number of intrusive memories. As predicted, prior to intervention there is no significant difference in intrusive memories between participant groups in the 24 hours following initial exposure to trauma stimuli (GLM: χ^2^=0.834, df=3, p=0.841), the mean number of intrusive memories is 3.334 (+/- 0.268). For the iM state (where p_2_+ p_3_=1; SI) the probability that intrusive memories reconsolidate unaltered (p_2_) is 1-exp(−3.33)=0.964 and hence the probability of reactivation (p_3_) is exp(−3.33)=0.036.

From the diaries, the mean number of intrusive memories over the whole week are 5.111 (+/- 0.996) for the no task control group, 1.889 (+/- 0.411) for the memory reminder cue+task group, 3.83 (+/- 0.682) for the task only group and, 4.889 (+/- 0.828) for the memory reminder cue only group. Again, using the discrete-valued Poisson distribution approach, the memory reminder cue group allows the probability that reactivated memory remains reactivated (p_4_) to be estimated, as this was the group to receive only the memory reminder cue. So, from this group the number of intrusive memories reflects memories in the reactivated state and the probability of no intrusive memories is 1-exp(−4.889)=0.9924. However, this probability combines p_3_ and p_4_ (as this group only had the reminder cue then recorded intrusive memories), so the marginal probability for p_4_ is the product of this joint probability and the probability of memory reactivation (p_4_= p_3_ (p_4_ ⋂ p_3_)=0.035).

Similarly, from the memory reminder cue +task group, the probability that memories successfully reconsolidate into the non-intrusive memory state (p_6_) via the treatment intervention can be determined. From this group, the probability of no intrusive memories is exp(−1.889)=0.151. Again, this is a combined probability of a reminder cue and a reconsolidation process (p_3_ and p_6_) so the marginal probability for p_6_ is the product of this joint probability and the probability of memory reactivation (p_6_=p_3_ (p_6_ ⋂ p_3_)=0.005). Using information from the reactivated memory state that p_4_+ p_5_+ p_6_ =1 (SI), the probability that a reactivated memory reconsolidates as an intrusive memory (p_5_) is simply determined from 1 – p_4_ – p_6_=0.959.

Using these transition probabilities, the stochastic model predicts low/intermediate persistence of iMs; this is principally driven by a combination of a high probability of intrusive memory reactivation with a high probability of intrusive memories reconsolidating in an unaltered way (Figure 2a). Critically, this analysis suggests that while the task is effective, maintaining intrusive memories in a reactivated state (p_4_) is essential to allowing non-intrusive forms of the memory to be reconsolidated (SI). The predicted long time to reach the non-intrusive memory state is constrained by a physical limit (Figure 2b) preventing opportunities for further effective task interventions.

**Figure 2.**
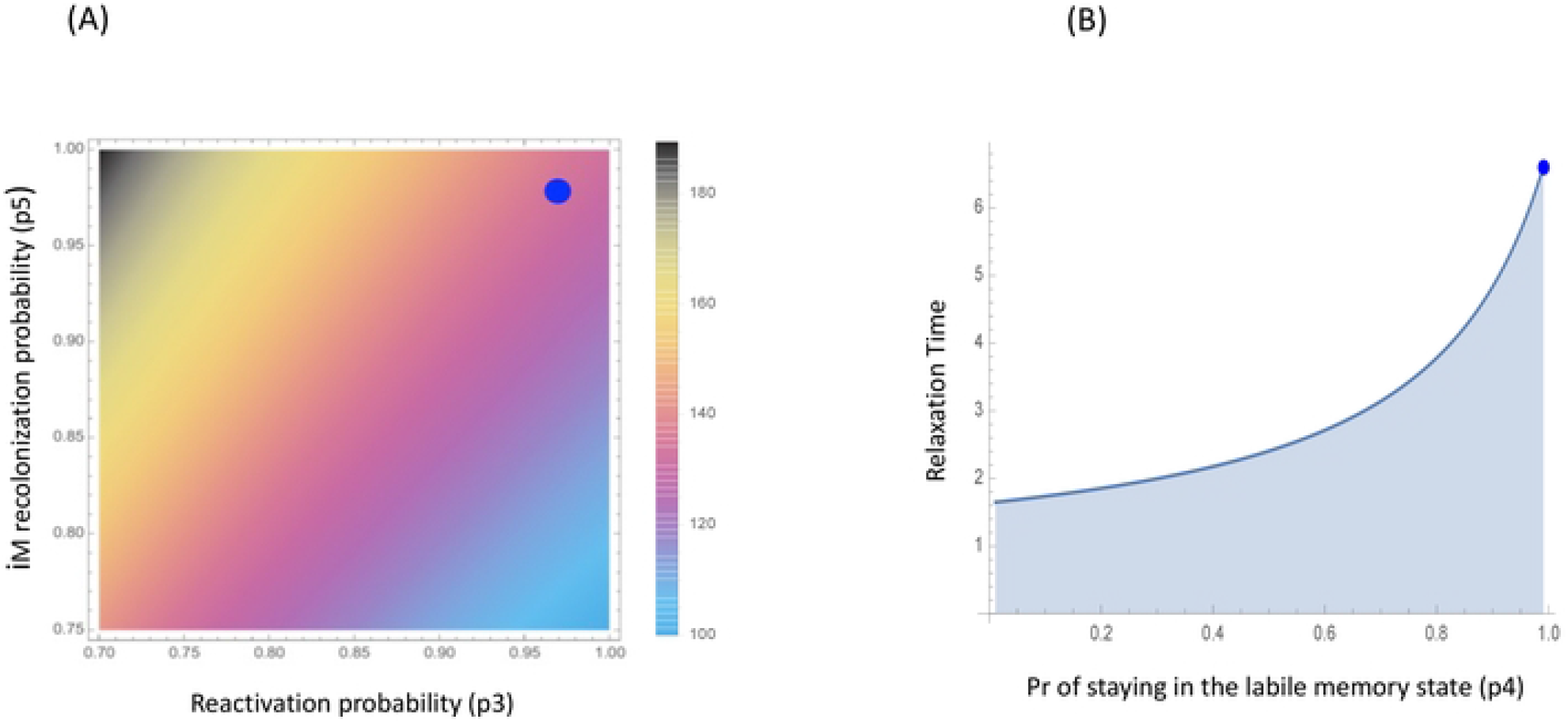
Stochastic trauma model predictions. Using the experimental data [29] analysis shows expected time in the intrusive memory (iM) state increases as reconsolidation probability (p_5_) increases and reactivation probability (p_3_) decreases. From the empirical parameterization of the unknown transition probabilities (p_2_ through to p_6_), the stochastic trauma model predicts (A) high reactivation and recolonization probabilities (blue dot) leading to intrusive memories that have low to intermediate persistence times in the consolidated iM state. (B) Time for memories to transit (so-called relaxation time) into the non-intrusive form of memory (niM) state have a physical limit (solid line) and for the experiments this time is expected to be low (blue dot).

### Simulating different treatment interventions

By definition, intrusive memories are those which come to mind involuntarily. The number of times a memory intrudes can be counted and recorded (say, in a diary). A reduction in the probability of the number of intrusions over a given time period is a primary outcome measure for recent intervention development [30,31,32]. Our stochastic framework can be used to simulate different treatment interventions. The expectation is that task memories (memories that are encoded during an intervention) interact and interfere with intrusive memories, for example, by competition for limited cognitive resources. By deriving a time-inhomogeneous version of the stochastic model (SI), different combinations of intervention components can be investigated. Delivering a single dose of task in the first time period, allows us to evaluate the long-term probability of memories successfully being rendered non-intrusive or returning to an intrusive memory state (Figure 3a). Increasing task strengths decrease the probability of intrusive memories remaining unaltered.

**Figure 3.**
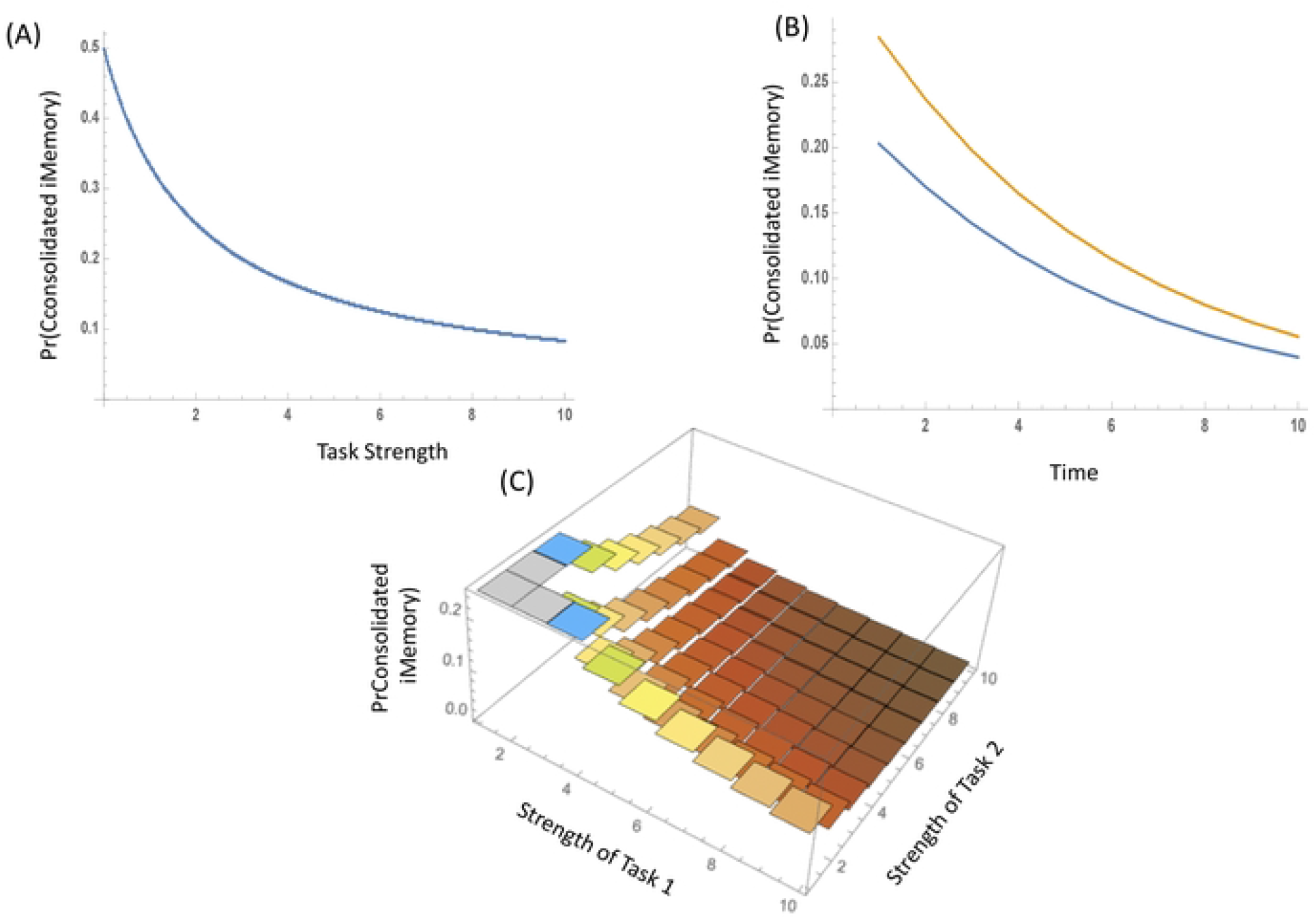
Simulation outcomes of trauma memory model for the effects of task strength on the probability of intrusive memories (iM). (A) hypothesis: how does task strength affect the probability of reconsolidated intrusive memory by delivering one dose of a task (in the first time period)? Simulations reveal that the probability of intrusive memories declines for increasing task strengths. (B) hypothesis: what is the role of task on the probability of intrusive memory reconsolidation over time by delivering one dose of task (in the first time period)? Simulations show that a task that interferes with the intrusive memory (blue line) is more likely to reduce the probability that intrusive memories reconsolidate compared to no task (orange line). (C) hypothesis: what is the effect of multiple tasks on the probability of intrusive memory reconsolidation? Simulations reveal that combining more than one task (in the first time point), of certain task strengths leads to stronger reduction in intrusive memories reconsolidating.

Delivering a single dose of task in the first time period has greater marginal gains in reducing the probability of intrusive memories reduction than no task interventions (Figure 3b). However, over time, these differences reduce and altering the task or task parameters might be necessary to prevent the intrusive memory reoccurring. Combining multiple tasks (say, two types of behavioural tasks) that act synergistically (additively or multiplicatively) can have greater effect at further reducing the probability of intrusive memories reoccurring. Delivering multiple doses of task(s) in the first time period is expected to achieve greater reductions in patterns of intrusive memories occurring than single tasks (Figure 3c).

Multiple independent memory reminder cue events (p_3_^n^; where n is the number of reminder cue events) interact with task strength to affect the probability of intrusive memories. Delivering multiple reminder cues under different task intervention can affect intrusive memory reoccurrence (Figure 3a-b). Critically, under weak task interventions, multiple reminder cues can increase the likelihood of intrusive memories reoccurring (Figure 4a-b) and thus worsen symptoms. Coupled with high probability of intrusive memories reconsolidating into their original form (p_5_), multiple reminder cues acting independently reduce the probability of reactivation (p_3_) and increase intrusive memory reoccurrence. In contrast, with conditionally-dependent reminder events (whereby the strength of subsequent reminder cues weakens compared to the strength of the previous reminder cue) then there is no interaction between task intervention and reminder cue; task interventions act to reduce the probability of intrusive memories reoccurring (Figure 4c-d) and may improve symptoms.

**Figure 4.**
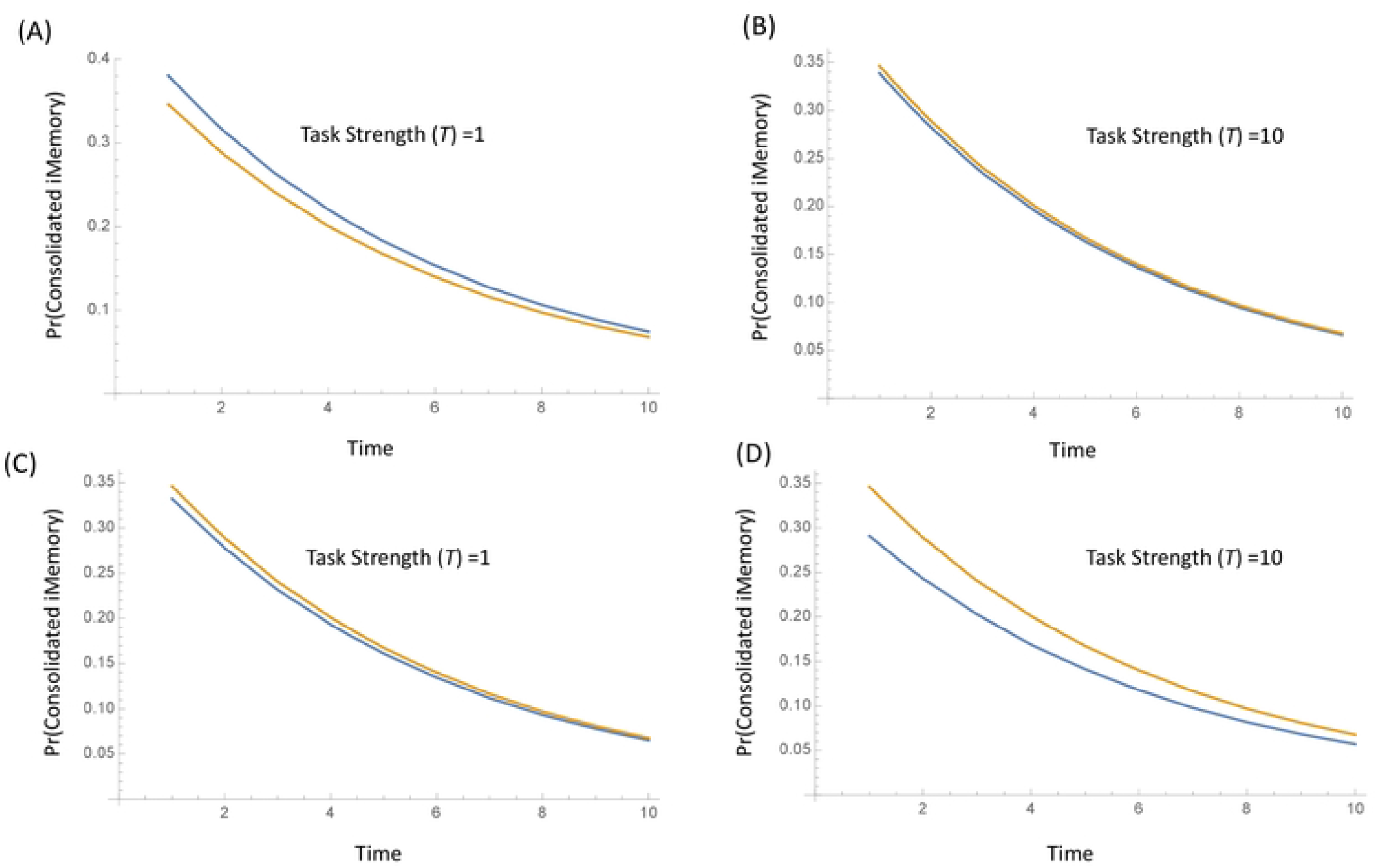
Simulation outcomes of trauma memory model for the effects of reminder cue on the probability of intrusive memories (iM) reconsolidating. (A-B) Outcome on the probability that iMs reconsolidate vary with multiple independent reminder cues and task strengths. (A) When the task strength is low (T=1), multiple reminder cues increase likelihood of intrusive memories reconsolidating (task blue line; no-task orange line). (B) When task strength is high (T=10), multiple reminder cues interact to affect the efficacy of the task (no difference between task/no task outcomes). Multiple reminders weaken task interventions in prevent iMs reconsolidating. (C-D) In contrast, under conditionally-dependent reminder cues (where the strength of cue weaken compared to the magnitude of the previous cue), then there is no interaction between task and reminder cue. Reminder cue together with the intervention task can reduce the probability of intrusive memories reconsolidating (task blue line; notask orange line).

## Discussion

Here, we have introduced a quantitative framework for understanding the modification of intrusive memories after traumatic events, and how targeting them in an intervention may help make them become less intrusive. We show how coupling reminder cues and task strengths can influenced the likelihood of reducing the reoccurrence of intrusive memories. We show, empirically, how the model framework can be used to evaluate the success of interventions and the key model sensitivities that allow intrusive memories to persist. Critical to this, the model framework develop here provides a predictive approach to understanding components of treatment interventions (e.g. task doses, reminder cue frequency) which have important clinical implications.

### Memory reactivation strength and frequency matters

Maintenance of different memories is affected by reactivation cues (e.g., [7]). For example, single presentations of a conditioned stimulus can induce reconsolidation and influence memory persistence. However, multiple cues can disrupt memories and lead to loss of acquired conditioned responses [7]. Here, in this study, we have shown that the number of repeated memory reminder cues affects memory persistence and multiple independent reactivation cues can render iMs *more* intrusive (for example multiple reminder cues can weaken task interventions, Figure 4). By contrast, *weakening* multiple reactivation cues can reduce the probability that reactivated, labile iMs reconsolidate.

Further to understanding memory reactivation and memory lability is how the strength of the cues can weaken or strengthen a memory. For instance, moderate levels of memory (re)activation are argued to be sufficient to lead to forgetting a memory [33]. Under a no-think/think paradigm, a non-monotonic relationship exists between memory activation and the consequential strength of the memory [33]: weak activation has limited effect on weakening a memory; moderate activation has optimal effect of memory weakening; while strong activation can strengthen the memory. Moreover, *incomplete* reminder cues which lead to prediction errors (differences between prior learned experience and a contemporary reality) allow memories to be destabilized, become labile and modified [34]. Here, we find that *weakening* sequential reminder cues can reduce intrusive memories: further investigating how pre-existing expectations, the type of the reminder cue and intrusive memory reactivation lead to new learning, memory encoding and reinforcement necessitates future study.

A corollary of all this is that intrusive memories operate within networks of brain architecture – changes in the amygdala, hippocampus and pre-frontal cortex occur following traumatic events [35]. Using real-time neural measures allows loops and networks across brain activity to visualized [36]. So, if the strength or number of reactivation cues lead to non-linear patterns in the changes to the reconsolidation of an intrusive memory and/or its suppression by reconsolidation of a neutral memory then further study, extending the Markov chain framework we develop here, is clearly warranted. Network-level effects of competition between iMs and niMs, the disruption of intrusive memory reconsolidation across an emotional-memory network and how information on consolidation/reconsolidation flows through these sorts of networks are all amenable questions within the stochastic modelling framework developed here.

Our framework suggests that briefer memory reactivation cue durations without multiple repetitions would be preferable for treatment success. This is of key clinical interest as current evidence-based psychological treatments [37] involve deliberately recalling the trauma memory often in a prolonged (and repeated) way, which while a form of treatment in itself, can be aversive and lead to patient drop out [38,39]. Shortening the duration of the memory reactivation cue may not only help make treatment more effective but also more tolerable for patients and could increase successful completion rates in therapy. The quality of memory reminder cues to achieve memory reactivation and adaptive memory updating requires calibration and may draw on insights from non-trauma memory [40].

Furthermore, our framework suggests increasing the strength of the task procedure is associated with poorer outcomes (see Figure 3). Many of those delivering clinical treatments and/or support after trauma might assume that conducting a longer and more intense treatment procedure(s) (here modelled as task strength) would be better than shorter ones. Our results suggest the reverse: decreasing the strength task procedure is associated with more beneficial outcomes in reducing the number of intrusive memories. Overall, this opens the intriguing possibility of optimising mental health treatments via research-driven insights from a mechanistically-driven, quantitative framework, rather than relying solely on practice-driven conventions that continue to dominate mental health research. To eliminate the recurrence of intrusive memories it may be optimal to use briefer and more focussed procedures targeting one intrusive memory at a time, rather than long and intense sessions reliving a whole trauma episode.

### Task interventions can influence suites of memory states

Following memory reactivation, both pharmacological and non-pharmacological interventions can interfere with memories. Studies have shown how different interventions influence memory states [7]. Here we show that interventions, tacitly through non-pharmacological approaches [29, 41], can determine times memories are in an intrusive state and as task strength increases, the time before memories enter a neutral state. For many people, intrusive memories following trauma might weaken over time without intervention [42–44]. However, for some they do not, so these sorts of interventions can be highly beneficial. Laboratory and clinical studies have shown that treatment interventions with a cognitive task can reduce the propensity of the intrusive memories to (re)consolidate following a memory reminder cue soon after a trauma [29,30,31,45]. Furthermore, there is emerging evidence for the success of these intervention when delivered a later time intervals since the trauma occurred [32,46,47]. Here, we have shown that delivering multiple doses or different (task) interventions is likely to achieve greater marginal gains in reducing the probability of intrusive memories reconsolidating than simply increasing the strength of a single task intervention.

### Bounds on outcomes

General and empirically-derived predictions from the stochastic framework highlight that there are bounds on memory reconsolidation outcomes following reactivation. Different combinations of task intervention strength and reactivation cue strength can lead to the same outcome in minimizing the time before memories consolidate into a neutral state. However, bounds exist on the time taken for memories to enter this state and these are critically dependent on the length of the reconsolidation window. Maintaining, with high probability, memories in a labile state, can lead to longer times for memories to consolidate into a neutral state. Low probability of maintaining memories in this labile state can limit time available for modification of these emotional memories. Understanding the critical physical time constraints on optimizing outcomes may require incorporating the details of neural circuitry dynamics (to understanding how neurons inhibit and excite to influence the length of the reconsolidation window) together with the time required to interrupt intrusive emotional memories with competing tasks.

### Framework for testing cognitive models of traumatic memory

Here, we introduce a model framework that is distinct in that in provides a conceptual way to synthesize the process of memory consolidation and reconsolidation. It is amenable to direct parameterization from experiments and has the value to be use as a part of clinical tools for the assessment and evaluation of interventions aimed at reducing the persistence of intrusive memories after traumatic events.

A unique advantage of our quantitative framework is that it links cognitive perspectives of trauma to the processes of memory reconsolidation. While cognitive conceptual models for the implications of trauma are well developed [20–23], they lack the mechanistic detail we develop here. These cognitive conceptual models are underpinned by the memory of the trauma being characterised by the frequency of involuntary intrusive memories. Early social-cognitive models such as Horowitz’s formulation of a stress-response syndrome [20] focus on the interplay between completion tendency (integrating trauma information on acceptable cognitive world model) and psychological defenses to keep the trauma information in an unconscious state; it is then this oscillation between integrating trauma and psychological defenses that lead to flashbacks and intrusions. Critically Horowitz’s formulation emphasizes the dynamic nature of trauma memory consolidation. Through our framework, this cognitive model is directly amenable to testing through an understanding of the probability by which memories consolidate - here we have assumed that trauma always leads to intrusive memory consolidate (p_1_ à 1.0). However, that need not be the case and building more dynamic, information-processing structures into the consolidation of trauma memory will allow different cognitive models of traumatic memory to be validated.

Under alternative conceptual frameworks, versions of the so-called dual representation theory [24, 48] posit that intrusive memories occur as an imbalance between the strengthening of emotion-laden sensory-bound representations and weakening of contextual representations in which the traumatic event occurred. Either strengthening of self-to-object (egocentric) and/or weakening of object-to-object (allocentric) memory processing can lead to the development of more intrusive memories [25, 49, 50]. The framework we develop here investigates the memory reconsolidation processes associated with changes in allocentric memory effects. Straightforward extensions of the mathematical framework, coupling different stochastic Markov chains, developed in this study, could allow versions of the dual representation theory to be parameterized. These coupled Markov chains could then allow predictions of both egocentric and allocentric memory processing of traumatic events to be compared and contrasted. Together with the mathematical approaches developed in our work here and elsewhere [26, 27], this may allow a way to combine mechanistic neural detail and cognitive process for greater understanding of mental health disorders.

### Conclusions

In conclusion, together with themes presented elsewhere [24,25], the stochastic modelling approach developed here provides a hierarchical, mechacognitive framework in which it is now feasible to embed neural mechanisms and cognitive processes. The fact that the stochastic framework opens up new ideas and provides a unique way in which to explore memory consolidation and reconsolidation is compelling for further developments [51]. Predictions that less reactivation strength and weaker task strengths favour a reduction in intrusive memories is intriguing. That traumatic events are limited to a small number of different intrusive memories, and that these forms of memory are amenable to intervention in different situations (road traffic accidents; traumatic childbirths; work-related trauma of intensive care unit staff), provides empirical support for mechanistically-driven, quantitative frameworks to extend our understanding of the temporal dynamic processes of treatments to reduce the persistence of intrusive memories after trauma.

## Methods

### Quantitative Framework

To model intrusive memory temporal dynamics we use a Markov chain approach. This aim of this framework is to capture the effects of an intervention (in our case a behavioural intervention; but the framework is equally applicable to pharmacological interventions) on intrusive memory (re)occurrence. Using this probabilistic model, memory states can be described as sequence of events in which the probability of transiting between states only depends on the state of the system at the previous event point. For modelling intrusive memories, we divide the Markov chain into four states: (i) prior trauma, no intrusive memory, (ii) a consolidated intrusive memory state, (iii) a reactivated intrusive memory and (iv) a non-intrusive memory state (see Figure 1a). In matrix form this is represented by:

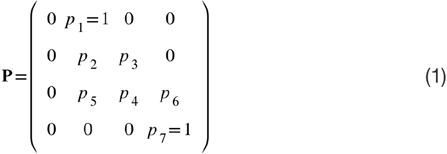

where p_i_ is the transition probability for the i^th^ event. p_1_ is the probability that an intrusive memory consolidates and is laid down as a memory. For this version of our model we assume that this always occurs. p_2_ is the probability that an intrusive memory reconsolidates and p_3_ is the probability that the intrusive memory is reactivated. p_4_ is the probability that reactivated memory stays reactivated, p_5_ is the probability that a reactivated memory reconsolidates as an intrusive memory and p_6_ is the probability that the reactivated memory reconsolidates as a non-intrusive memory (niM). p_7_ is the probability that a non-intrusive memory (niM) remains in this state (here, we assume this in absorbing state so p_7_=1.0). Probabilities in each of rows of the Markov chain sum to 1.

### Model Parameters

Our aim is to understand how a behavioural task intervention and/or a reminder cue affect the probability of intrusive memories reconsolidating after reactivation into a non intrusive state. This task intervention is described by its effects on the probability of a reactivated iM reconsolidating into the iM state. In the matrix (eqn 1), this is probability transition p_5_ and, in a general form, we model this sort of task intervention as:

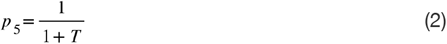

where *T* is the strength of the task intervention. As *T* increases the task intervention is more effective, p_5_ decreases and p_6_ increases (p_6_ = 1 – p_4_ – p_5_). We consider the role this task has on influencing probabilistic outcomes of memory reconsolidation. To describe the probability of reactivation (p_3_) following a reminder cue, we assume that this can be derived from a binary process and as such probabilistically can be represented, in general form, as a logistic function:

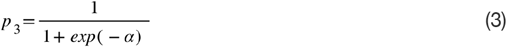

where α is the strength of the reminder cue.

### Analysis

Using this stochastic approach to model the (re)consolidation of intrusive memories, it is feasible to determine measures such as (i) the expected time to absorption into the non-intrusive memory (niM) state, (ii) number of visits to the reactivated iM state before absorption into the niM state and (iii) how long memories stay ‘mixed’ in different states - all as a function of task and/or reminder cue strength. This is achieved through analysis of the Markov chain (see below) as the characteristic polynomial of a Markov chain (from Det(**P** - λ**I**) allows eigenvalues and (right) eigenvectors (**V**) to be determined. Using spectral decomposition yields an expression for the long-term probabilities of memory states: **P**^n^ = **V D**^n^**V**^−1^ where **D** is a diagonal matrix of eigenvalues. We develop this approach to analyse the temporal dynamics of intrusive memory reconsolidation and further details on the analysis and numerics are given below.

## Long-term temporal states

To explore the efficacy of task interventions at redistributing the probability of memories being in different states (consolidated iM, reactivated iM, reconsolidated non-intrusive memory - niM) we determine the long-term state probabilities. For transition matrices where probabilities are fixed (e.g., interventions operate at each time step, or there are no interventions) across all time, then:

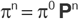

where π^0^ is the initial state vector (the distribution of memories in different starting states), **P** is the time-homogeneous transition matrix (eqn 1) and π^n^ is the state vector after n time steps. The long-term state of the system is determined from **P**^n^ = **V D**^n^**V**^−1^.

For transition matrices where there are differences between the first and subsequent transitions (time-inhomogeneous chains) such that only the first transition includes the reminder cue and task interventions (affecting transition probabilities) then the long-term evolution of the probabilities of being in different memory states (π) at the n^th^ time step is given by:

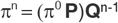

where π^0^ is the initial state vector, **P** is the initial transition matrix that includes the reminder cue and task intervention, and **Q** is transition matrix that does not include the reminder cue or the task intervention conditional probabilities. Spectral decomposition is then a two-step process. The first transition is determined from:

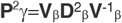

where **V**_β_ are eigenvectors and **D** is a diagonal matrix of eigenvalues. The subsequent transitions are:

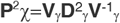

where **V**_γ_ are the eigenvectors and **D** is the diagonal matrix of eigenvalues determined from the characteristic polynomial of **P**^2^γ=Det(**P**γ - λ**I**) (where λ are eigenvalues).

## Time to Absorption

### Absorbing Markov chains with limiting transition matrices

Given that the chain is in a particular memory state, a key question to ask is: what is the expected number of steps before the chain is absorbed into the non-intrusive memory (niM) state? For the absorbing Markov chain the matrix **I**-**P** has an inverse **N** and **N** = **I** + **P** + **P**^2^ + …. The ij-entry n_ij_ of the matrix **N** is the expected number of times the chain is in the j^th^ memory state given that the chain starts in the i^th^ memory state.

### Absorbing Markov chains with varying transition matrices

As noted above, with a time-homogeneous transition matrix:

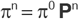

holds for determining the long-term state of system under strong ergodicity. For time-inhomogeneous Markov chains, weak ergodicity is expected and:

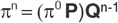

where **P** and **Q** are different transition matrices. A ***weak ergodic condition*** states:

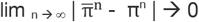

where 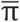 and π are different state vectors. So, a trivial generalisation of the strong ergodic condition is then:

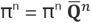

where 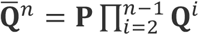. This weak ergodic condition also implies that 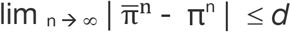 (where *d is* some minimal difference) approximates the strong ergodic condition, and the application of these conditions to time-inhomogeneous matrices has been investigated [52, 53].

### Canonical Form

Given this weak ergodic condition, as noted, the transition matrices can then be expressed as:

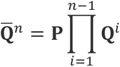

So, the time-inhomogeneous transition matrix can be expressed in canonical form [54]. If q_ij_^n^ of transition matrix 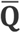 is the probability of starting in state *s*_i_ and being in state *s*_j_, after n time steps then the canonical form (following [54]) is:

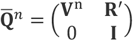

where V^n^ is a matrix of eigenvectors at the n^th^ time point, **R**′ is absorbing state matrix and **I** is an identity matrix. This allows an expression for time in states for Markov chains with varying transition probabilities to be used (see below).

### Time in states

Let q_ij_^n^ be the entry in **P**^n^. Let *Z*^t^ be a random variable equal to 1 if the chain is in state *s*_j_ after t time steps or 0 otherwise. Then:

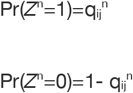

The expected time in state is then:

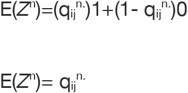

Across all transient states, *s*_j_, then:

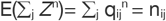

### Spectral gap and mixing times

Eigenvalue properties can be used to determine how long states stay ‘mixed’. Eigenvalues for transition matrices are often labelled:

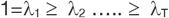

and the largest eigenvalue with absolute value | λ*| is defined as:

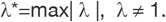

The spectral gap (defined as the difference between the moduli of the two dominant eigenvalues in a transition matrix) is γ = 1- λ* and the relaxation time to convergence is τ =1/ γ. These eigenvalue properties can be used to determine how long different memories stay ‘mixed’

A further problem is ‘how fast is convergence to the non-intrusive memory state?’ or as a corollary, what is an upper bound on the convergence of the Markov chain to a fixed steady state. That is:

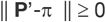

This requires the definition of an appropriate distance metric and here we chose a *total variation distance* (TV) where:

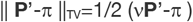

where v is a scaling parameter. So

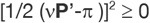

Expanding this expression yields:

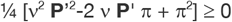

Setting a= **P**’^2^, b=2 v **P**’ π, c= π^2^, and v =b/2a, yields:

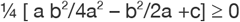

So,

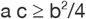

This inequality can be rewritten as:

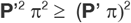

The upper bound on convergence to the initial state is a Cauchy-Schwarz inequality [55] such that || **P**’-π || ≥ 0 if

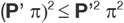

We use this analysis and bound to understand the rate at which intrusive memories consolidate into a neutral state (see Figure 2B).

## Numerics

For then numerical analysis and to investigate model predictions, we use the following formulations of the Markov chains and a canonical set of parameters.

To investigate persistence times (figures 1b-d), we use the following set of transition probabilities and parameter values: strength of reminder cue α=1.0 and probability of reactivated memory staying reactivated p_4_=0.5.

To investigate persistence times (figure 2), we use the following set of transition matrices:

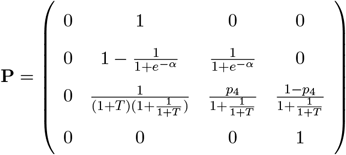

where strength of reminder cue α = 0.5 and probability of memory staying reactivated p_4_=0.5, and

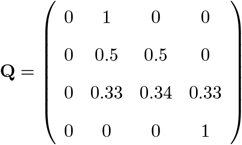

The initial memory state vector was π=[0,0.5,0.5,0]^⊤^. For the two tasks (T_1_, T_2_) acting multiplicatively, we use:

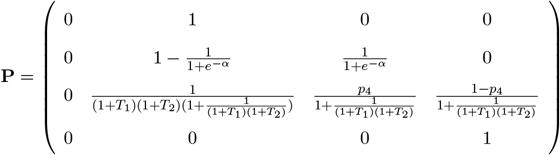

with strength of reminder cue α = 0.5 and probability of reactivated memory staying reactivated p_4_=0.5. Initial memory states were π=[0,0.5,0.5,0]^⊤^.

To investigate the effects of the reminder cue (figure 3), we use the transition matrices as for figure 2 (given above) with p_3_=(1/(1+exp(−α)))^5^ to investigate the multiplicative effects of the reminder cue. For the conditional-dependent reminder cue, we use p_3_=1/(1+exp(−p_3_’’)) with p_3_’’ =1/(1+exp(−p_3_’)) and p_3_ ‘ =1/(1+exp(−a)) with α = 0.5 and p_4_=0.5. Initial memory states were π=[0,0.5,0.5,0]^⊤^.

All analyses were completed in Mathematica and the scripts are available at the Open Science Framework: \url{https://osf.io/v4ynf/}.

## Acknowledgements

We thank Philip Millroth and Renée Visser for constructive comments on this work. EAH receives funding from the Swedish Research Council (2020-00873), The Oak Foundation (OCAY-18-442) and The Wellcome Trust (223016/Z/21/Z).

## List of supplementary materials

1. Worksheet determining transition probabilities from designed experimental data

